# Mapping global land sharing-sparing patterns between human and wildlife

**DOI:** 10.1101/2022.09.09.507273

**Authors:** Chengcheng Zhang, Yihong Wang, Shengkai Pan, Biao Yang, Xiangjiang Zhan, Jiang Chang, Junsheng Li, Qiang Dai

## Abstract

Understanding the global patterns of land sharing-sparing between humans and wildlife is essential for pragmatic conservation implementation, yet analytical foundations and indicator-based assessments are still lacking. By integrating distributions of 30,664 terrestrial vertebrates and human pressures, we provide a series of spatial explicit Human-Nature Indices (HNIs) before classifying the global lands into four categories. We found that the *Co-occurring* (*C*) regions, where lands are shared by humans and wildlife, are not insignificant (16.91% of global land). For land-sparing, the *Diversity-intact* (*D*) and the *Anthropic* (*A*) regions account for 45.64% and 1.41% of the land, respectively. The patterns of HNIs, varying considerably among taxonomic groups, are determined mainly by the expansion of human-dominated land use. Land sharing and sparing could work as complementary strategies to support biodiversity and human development toward ambitious and pragmatic 30 by 30 goals. Our results highlight that those regions should adopt different conservation strategies according to their sharing-sparing patterns and distribution of protected areas.

## 1. Introduction

The human-nature relationship has always been complicated and ambiguous, nature being seen as both friend and foe (Bourdeau, 2004). Although Earth has been described as a stable and self-regulating life-environment system (Free and Barton, 2007), the ecosystems constitute the ‘life-support system’ for humans (Hayward, 1997). Nonetheless, human activities have been dramatically altering the land surface of our planet and threatening this system (McKee et al., 2004; Syphard et al., 2009). We live amid a global wave of anthropogenically driven biodiversity loss (Barnosky et al., 2011; Ceballos et al., 2015) which is so severe that the ‘people and nature’ framing has been becoming an emerging purpose underpinning conservation biology in recent decades (Mace, 2014) and biodiversity conservation has become a consensus among most countries (Chandra and Idrisova, 2011).

Among numerous human-induced environmental changes, land use change is believed to be one and the most significant influence exerted on the environment, caused by deep-rooted land use competition between humans and wildlife, over shelter and resources (Krausmann et al., 2013; McDonald et al., 2020). Urban land expansion, agriculture production, and livestock grazing have been escalating habitat loss, which is a leading cause behind biodiversity decline (Alkemade et al., 2013; McDonald et al., 2020; Norris, 2008; Sala et al., 2000). To reconcile the conflict between human activities and biodiversity conservation, the notion of ‘land sharing’ and ‘land sparing’ have been heated discussed (Crespin and Simonetti, 2019; Fischer et al., 2014; Salles et al., 2017).

Land-sharing and land-sparing conceptualize the spatial relationships between human and wildlife: land-sharing integrates both objectives on the same land, while land-sparing separates human activities from remnants of wilderness to avoid future human intervention. The land sharing-sparing framework was originally developed in the context of farming, in an attempt to meet rising food demand at the least cost to biodiversity (Green et al., 2005; Kremen, 2015; Phalan et al., 2011). Then, this framework has been put forward to provide potential solutions to biodiversity losses caused by urban development or ecosystem services (Gilroy et al., 2014; Ibáñez - Álamo et al., 2020; Lin and Fuller, 2013).

Protecting biodiversity while sustaining human development is one of our most significant modern challenges. Various ambitious and responsible solutions have been proposed to halt continuing biodiversity loss and plan practical conservation strategies (Immovilli and Kok, 2020; Maxwell et al., 2020; Pimm et al., 2018). Meanwhile, there is increasing and justifiable pressure to properly account for human needs when setting conservation goals (Dudley, 2008). Insufficient considerations of human needs often limit the further implementation of conservation planning and actions. From the perspective of the Convention on Biological Diversity, realistic concerns are of equal importance (CBD, 2018). As consideration has already been given to the Post-2020 Global Biodiversity Framework for conserving at least 30% of the Earth’s terrestrial lands, it is necessary that the parties take responsibility based on the actual situations and scientific assessments. However, without analytical foundations to inform the global land sharing-sparing patterns between human and wildlife and indicator-based assessment, it remains an obstacle to convert global targets into local ones for practical actions and guarantee the overall 30 by 30 goals. There is an urgent need to make the ‘people and nature’ framing measurable and facilitate its implementation.

Here we propose a series of spatial explicit Human-Nature Indices (HNIs) comprising four indices – *Anthropic* (*A*), *Barren* (*B*), *Co-occurring* (*C*), and *Diversity-intact* (*D*) *Indices* (Materials and Methods), to reveal the global land sharing-sparing patterns between human development and terrestrial vertebrate richness. Species diversity is derived from the refined distribution of 30,664 terrestrial vertebrate species, and human pressures are modified from the latest global human footprint map. The *D* index indicates the extent of biodiversity superiority over human activity in an area. The areas with high *D* values have high biodiversity and few human activities (Fig. 1). The *C* index represents the extent of co-occurrence between humans and biodiversity in a region, with high values indicating areas with both high biodiversity and intensive human activities. The *A* index indicates the extent that humans dominate lands. Where *A* index is high shows places with intensive human activities and low biodiversity. The *B* index shows the extent of how barren and desolate an area is, high values of which indicate both biodiversity and human activity are low in a region. The HNIs for the four taxonomic groups (amphibians, reptiles, birds and mammals) were also evaluated, separately. We then assessed the drivers of HNIs to understand how human development and environmental factors have been shaping the HNIs patterns. Finally, we present a global PA network assessment under the frame of the human-nature relationship. By classifying the global terrestrial lands into *A, B, C* and *D* regions before overlapping the PAs with the four regions, we assessed if the current PA network fulfilled the conservation requirement, and offered different strategies for different regions. We hope our framework can provide a new perspective for conservation implementation.

**Fig. 1.**
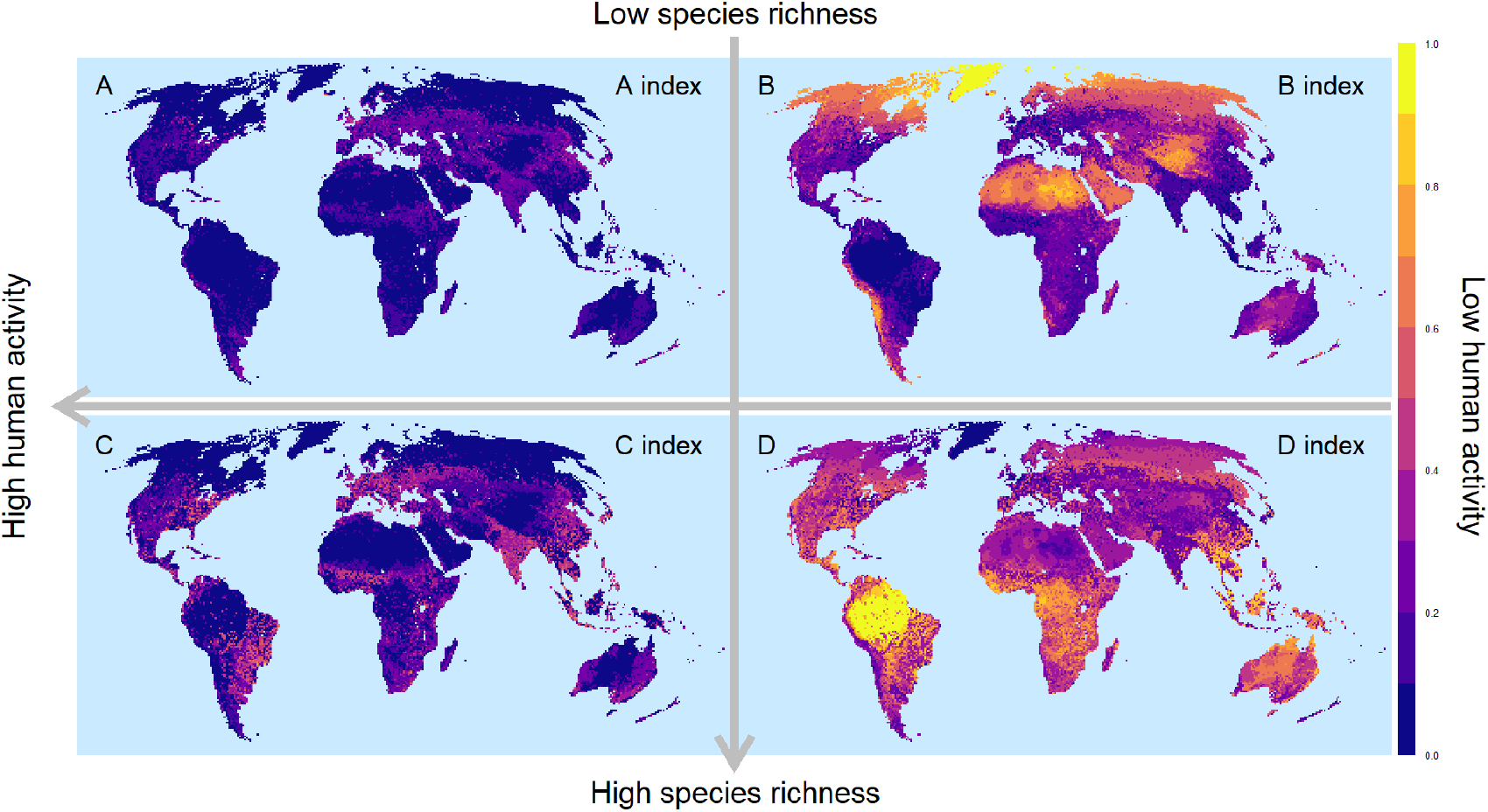
Global spatially explicit land-sharing and land-sparing patterns between human and terrestrial vertebrates illustrate by the four Human-Nature Indices (HNIs). (a) The *Anthropic* (*A*) index indicates the extent of humans dominating land; (b) the *Barren* (*B*) index shows the extent of how barren an area is for both human and wildlife; (c) the *Co-occurring* (*C*) index presents the extent of human and wildlife co-occurrence; and (d) the *Diversity-intact* (*D*) index indicates the extent that lands are dominated by wildlife.

## 2. Methods

### 2.1. Species richness

Global species distribution data were obtained from IUCN (version 2020-10) (IUCN, 2020) for 6803 amphibians, 7170 reptiles, and 5537 terrestrial mammals, while 11,154 birds distribution data were obtained from Birdlife International (version 2020.1) (BirdLife International and Handbook of the Birds of the World, 2020). The original data were available as species range polygons, which were reprojected into Mollweide equal area projection before being converted to raster layers with a cell resolution of 835m. Land sparing and sharing can show different patterns at larger scales, yet from conservation planning and implementation, the scale of about 1 km^2^ is plausible for PAs and other effective area-based conservation measures. A species was considered present in a raster cell if the cell intersects (contains, crosses, or within) the range polygon. We clipped the species distribution maps by suitable habitat for each species based on habitat type against the global terrestrial habitat types (Jung et al., 2020) and suitable elevation limits against the GMTED2010 elevation data (Danielson and Gesch, 2011). Species richness for each of the four taxonomic groups was calculated as the number of species present in each raster cell.

### 2.2. Human pressure

A modified global terrestrial human footprint (HFP) map was used to indicate the human pressure. The latest HFP (Venter et al., 2016) was referred to represent the most current information on both direct and indirect human pressures on the environment, which includes eight individual pressure layers: built environments, crop lands, pasture lands, population density, nightlights, railways, major roadways, and navigable waterways. Roadways and Built environments pressure layers were weighted by rescaled population density ([0, 1]), considering that constructions located in low population density regions would place less stress on the environment than that in high population density regions. Based on our field knowledge in the Qinghai-Tibetan Plateau and Xinjiang Autonomous Region, many areas belonging to pasture were misidentified as cropland. Thus, we used the cropland areas derived from ESA Climate Change Initiative Land Cover data (ESA, 2017) to correct them. The Population density data was not included in the human pressure map. This layer shows population density at a scale of administrative regions that does not represent its real impacts on ecological regions. Its influences are already indicated in the modified Roadways and Built environments layers. The weighted Roadways, weighted Built environments, modified Cropland, modified Pasture layers, along with the original Nightlights, Railways and Navigable waterways layers, were added together to form the human pressure map with a value range from 0 to 40. The values were log10 plus one transformed with range rescaled to [0,1] as the human pressure index.

### 2.3. Human-Nature Indices (HNIs)

We proposed four spatial indices - *Anthropic index* (*A*), *Barren Index* (*B*), *Co-occurring Index* (*C*) and *Diversity-intact Index (D)* to illustrate the global land sharing and land sparing patterns between human and wildlife:

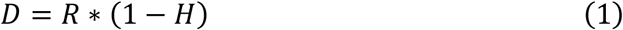

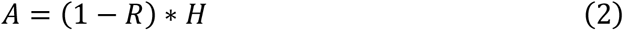

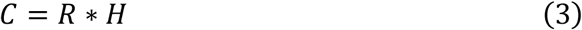

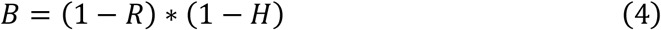

where *R* represents the biodiversity in a raster cell, *H* is the human pressure index. *D* index indicates the extent of a land dominated by species richness, while *A* index denotes how much land is dominated by humans. Together, they represent two opposite orientations of land-sparing. *C* index indicates the extent of land where species and humans co-occur, thus representing land-sharing. *B* index shows how rarely a land is used by wildlife or human beings.

We calculated those four HNIs for the four taxonomic species groups (amphibians, reptiles, birds and mammals) respectively and all species. For the spatial relationship between human and a given taxon, *R* is given by the following formulas:

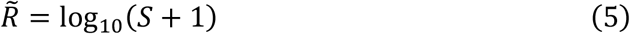

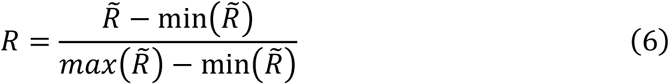

where *S* represents the species richness of the given taxon. When we synthetically assess the HNIs for all the four taxa, *R* is calculated as followings:

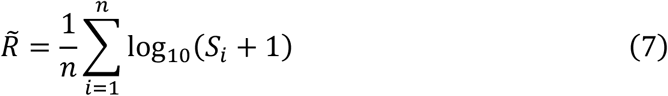

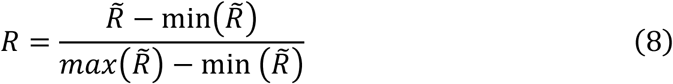

where *S*_*i*_ is the species richness of a given taxon, and n is the number of all taxa.

### 2.4. Environmental covariates

Four bioclimatic variables data (annual mean temperature, temperature seasonality, annual precipitation and precipitation seasonality) and elevation data at 30 arc-seconds resolution were obtained from WorldClim Version 2.1 (Fick and Hijmans, 2017). A measure of roughness, elevation standard deviation, derived from the MERIT-DEM data (Amatulli et al., 2020), which is in a resolution of 250m. A measure of long-term human activity, crop land percentage changes from 1700 to 1992 (Ramankutty and Foley, 1999) is available at a resolution of 5 arc-minutes. All covariates were reprojected to Mollweide equal-area projection and rescaled to 835m resolution before further analyses.

### 2.5. Statistical analysis

We used Random Forests to estimate the potential of each covariate in explaining the patterns of HNIs. Models were first fitted for the four indices for total species. Then we modeled the four indices to all the environmental covariates for the four taxonomic species groups separately. The permutation-based measure, per cent increase in mean squared error (%IncMSE), was applied to evaluate the variable importance. The correlation test among variables showed that only mean annual temperature and temperature seasonality are largely related, with an absolute correlation coefficient greater than 0.8. To explore the response relationships between each variable and the land sharing-sparing patterns, we computed the partial dependence function for regression then plotted their marginal effects. Models fitness and plots were carried out in R (version 4.0.3) with the packages of ‘randomForest’ (Liaw and Wiener, 2002) and ‘edarf’ (Jones and Linder, 2016).

### 2.6. Protected areas and gap analysis

Data on protected areas (PAs) were obtained from the September 2020 World Database of Protected Areas (UNEP-WCMC & IUCN) (UNEP-WCMC and IUCN, 2020) except China. For PAs in China, we used the dataset of the July 2017 WDPA, as it is more complete in that region. PAs were categorized into strictly PAs (IUCN I-IV categories) and multiple-use PAs (IUCN V-VI, not reported and not assigned categories). Many PAs in China with strict protection (He and Cliquet, 2020) fall into the category of V or VI. Therefore, for China’s PAs with a category of V or VI, we did not follow their IUCN statuses. Instead, we assigned them to the strictly Pas category when designated as ‘National Nature Reserve’ or ‘Nature Reserve’. We set them to the multiple-use PAs group when designated as ‘Scenic Area’. We tested the overlap and gaps between the global terrestrial PAs network and the *A, B, C* and *D* regions.

## 3. Results

### 3.1. The Human-Nature Indices (HNIs)

#### 3.1.1. Global patterns

For the relationship between total terrestrial vertebrates and human pressure, the regions with higher *Diversity-intact* values mainly aggregate in the Amazon-Orinoco Basin, the Congo Basin, and center and west of Australia, also scatter in southern North America, West Africa, East Africa, and Southeast Asia (Fig. 1d). Typical high *Co-occurring* value areas distribute in the east and center of United States, Mesoamerica and the northern Andes, the south and east of Brazil, forests and savannas of West Africa, the western and southern of Europe, South Asia, Southeast Asia and southeastern of China (Fig. 1c). In general, the areas that suit humans also suit plentiful wildlife at a global scale. Meanwhile, the high *Anthropic* value areas, characterized by intense human activity and low biodiversity, are rare across the globe (Fig. 1a). Despite the increasing expansion of human activities on this planet and their impacts on biodiversity, the areas merely dominated by humans are limited, and the *A* values are rarely higher than 0.5. The high *Barren* value areas are mainly distributed in the high latitude areas, the Qinghai-Tibet Plateau, the Arabian Peninsula, the Sahara Desert, and the middle Andes (Fig. 1b).

We then assess the HNIs for the four taxonomic groups (amphibians, reptiles, birds, and mammals). The four land-sharing and land-sparing patterns differ among taxa, especially between ectotherms and endotherms (Fig. S1-S4). From the two indices that account for terrestrial vertebrate richness, *D* and *C*, the areas with high *D* and high *C* values for birds and mammals are more extensive than those for amphibians and reptiles.

#### 3.1.2. Drivers of the HNIs

We used Random Forest to estimate the potential of each covariate in explaining the patterns of human-wildlife land sharing-sparing. On a global scale, the leading driver of this land sharing-sparing pattern is the expansion of human-dominated land use (Fig. 2). The *C* index showed a positive asymptotic relationship with long-term land-use change (Fig. 3a), while the *D* index showed negative (Fig. 3b). Mean temperature and annual precipitation also greatly influence these spatial patterns. The relationship between *C* values and mean temperature followed an S-shaped growth curve (Fig. 3e), while *C* values followed a hump-shaped response to annual precipitation (Fig. 3c).

**Fig. 2.**
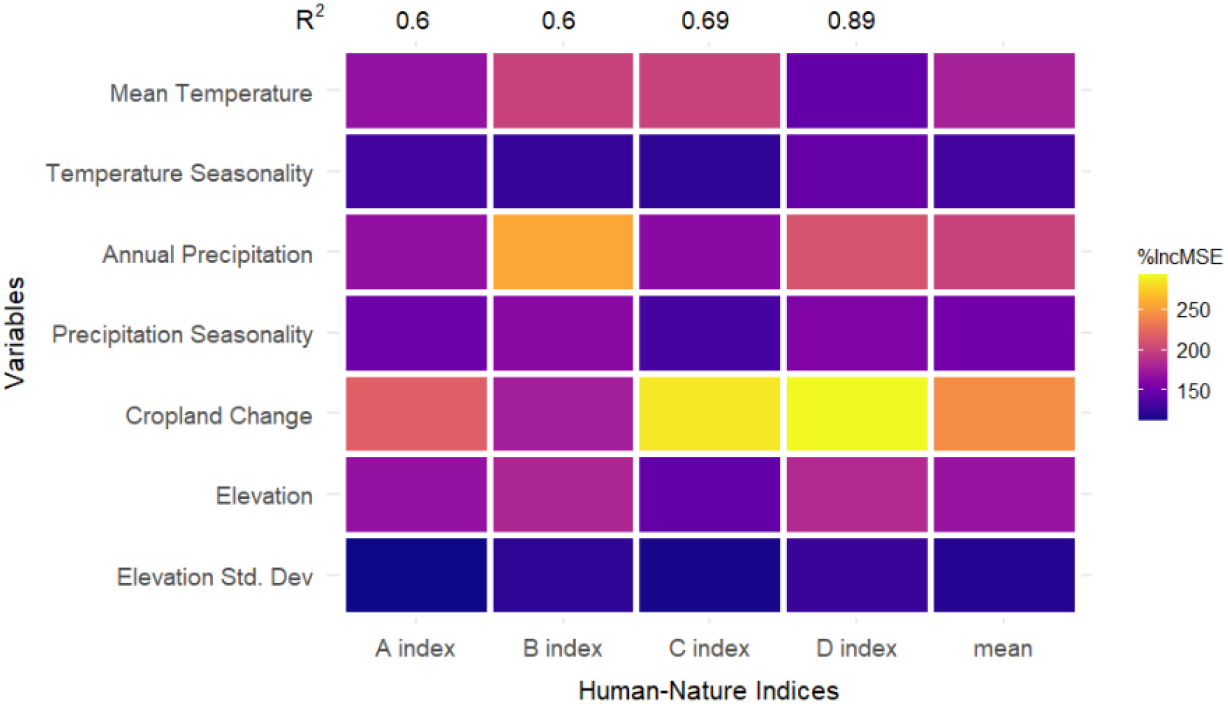
Importance of variables in predicting the four global Human-Nature Indices (HNIs) - *Anthropic* (*A*), *Barren* (*B*), *Co-occurring* (*C*), and *Diversity-intact* (*D*) indices. The per cent increase in mean squared error (%IncMSE) for each index was used to evaluate variable importance, indicated by the yellow (high importance) to blue (low importance) color gradient.

**Fig. 3.**
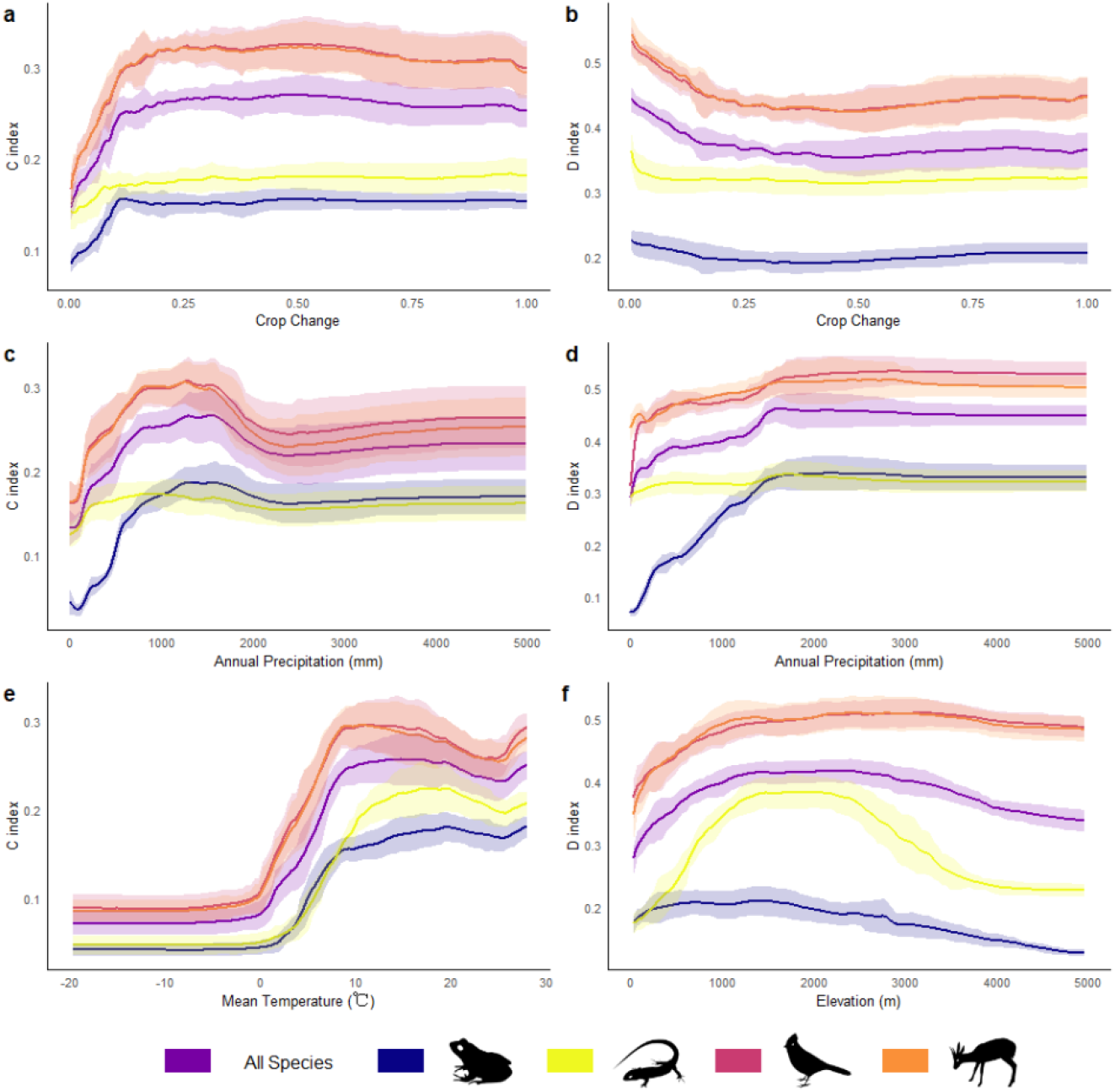
Relationships between a selection of the most important environmental variables and the *Co-occurring* index (*C* index, left column) or the *Diversity-intact* index (*D* index, right column). Line colors indicate different taxonomic groups: all species (purple), amphibians (blue), reptiles (yellow), birds (dark pink) and mammals (orange). Shaded areas indicate the standard deviation around the partial relationships.

### 3.2. The A, B, C and D regions

We then compared the values of the four HNIs in each grid cell. Where the value of the *D* index is greater than the values of the other three was considered as ‘*D* region’, indicating the areas that dominated by wildlife, resulting in a total area of 60.90 million km^2^ in terrestrial land (Fig. 4). Similarly, we defined ‘*C* region’ where human and wildlife co-occur (22.56 million km^2^), ‘*A* region’ that dominated by human (1.89 million km^2^) and ‘*B* region’ where both biodiversity and human activities is low (48.10 million km^2^). From the perspective of land sharing-sparing, the *C* region can be deemed as areas of sharing while the *A* and *D* regions as areas of sparing.

**Fig. 4.**
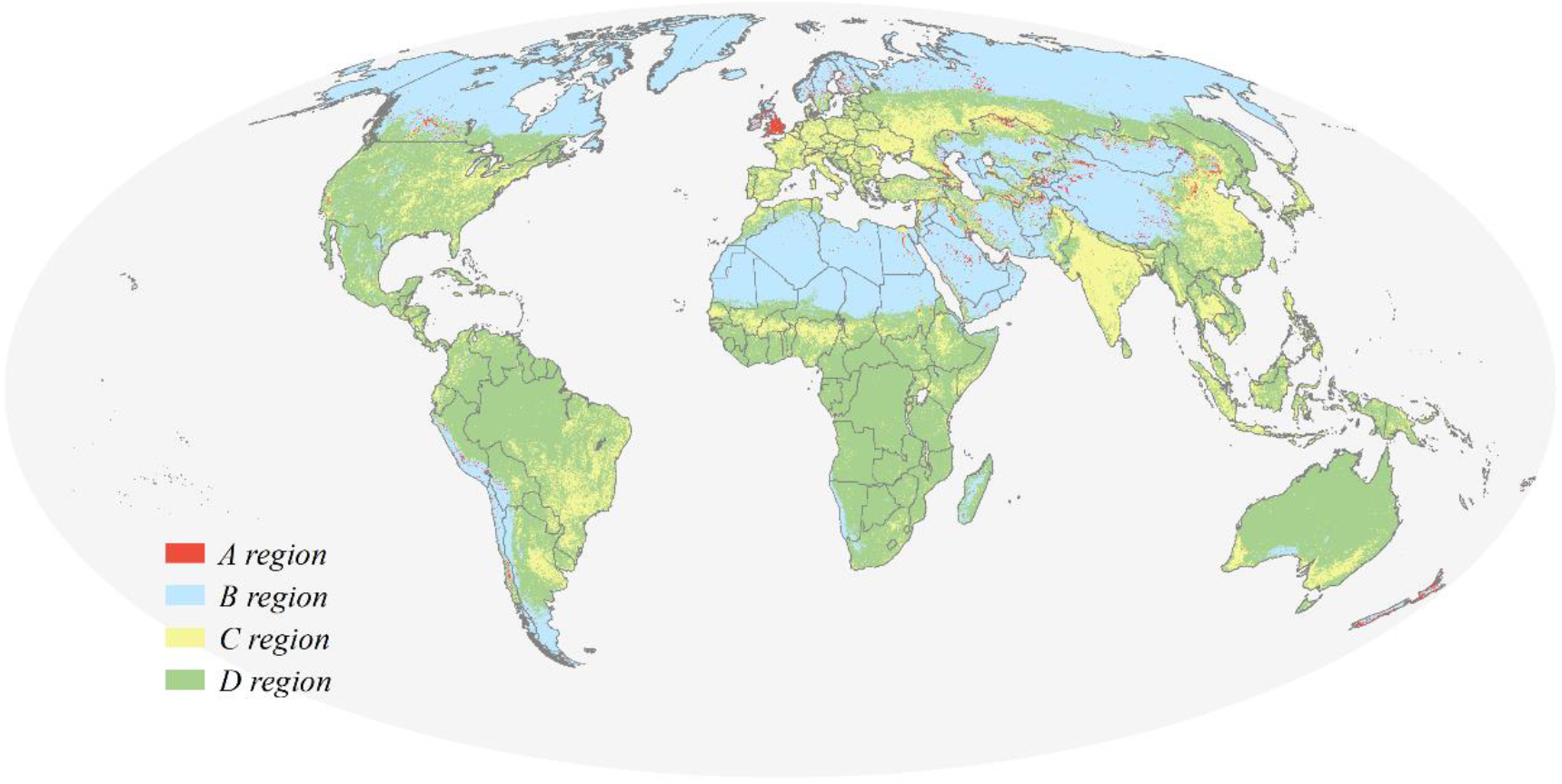
*Anthropic* (*A*), *Barren* (*B*), *Co-occurring* (*C*), and *Diversity-intact* (*D*) regions illustrate land-sharing and land-sparing statues between human and terrestrial vertebrates in different countries and territories.

#### 3.2.1. Variation between taxonomic groups

The divisions of the *A, B, C*, and *D* four regions differ substantially among the four vertebrate taxa (Fig. 5 & S5-S8). The distribution of *C* regions for birds and mammals (18.22% and 17.98% of land), extending from low-middle latitudes to high latitudes, is broader than those for amphibians and reptiles (5.14% and 8.44% of land). The land-sparing *A* and *D* regions show opposite trends between ectotherm and endotherm. In terms of area, the *A* regions of amphibians and reptiles take up 13.19% and 9.90% of global terrestrial land, respectively. They are mainly located in low and middle latitudes, including South Asia, East Asia, and Europe. In contrast to amphibians and reptiles, the *A* regions of birds and mammals are few, accounting for 0.12% and 0.36% of terrestrial lands. The *D* regions of birds and mammals have the widest distribution, accounting for 67.54% and 71.45% of the terrestrial land, respectively. However, Amphibians and reptiles have smaller *D* regions (21.15% and 27.25% of terrestrial land) that cover only low to middle latitudes. The *B* regions of amphibians and reptiles occupy large areas of terrestrial land (60.51% and 54.41%, separately). Whereas for birds and mammals, those are much less (14.12% and 10.22%, separately), locating in the Sahara, the Arabian Peninsula, Greenland, and other middle to high latitudes.

**Fig. 5.**
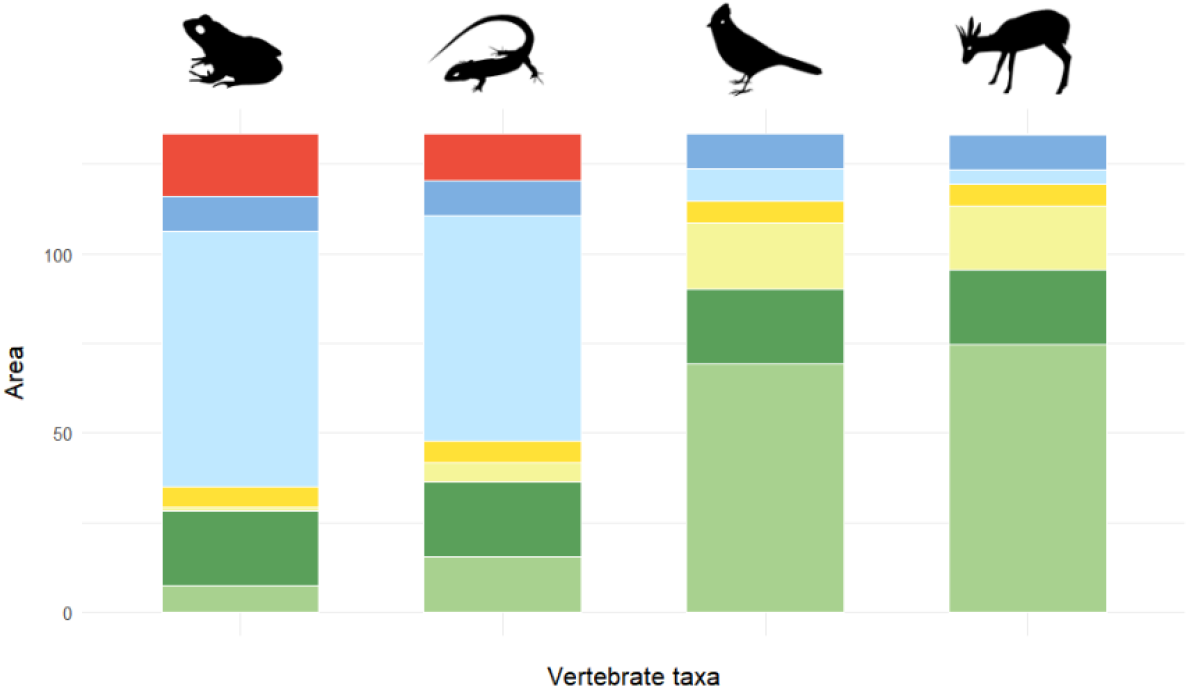
Partitions of the *Anthropic* (*A*), *Barren* (*B*), *Co-occurring* (*C*), and *Diversity-intact* (*D*) regions among the four taxonomic groups (amphibians, reptiles, birds and mammals). Red colors indicate the *A* region, blue colors indicate the *B* region, yellow colors indicate the *C* region, and green colors indicate the *D* region. Darker colors imply the areas shared by all the four taxonomic groups.

Fig. S9 maps the overlaps of the four regions of the four taxonomic group. Of which, 20.83 million km^2^, accounting for 15.59% of global land, are the *D* region shared by all the four taxa, mainly located in low to middle latitudes including South America, southeastern North America, West-Central Africa, Southeast Asia, and northeastern Australia. The overlapping areas of *C* regions usually surround those of *D* regions, and take up 4.51% of land area. The overlaps of *B* regions, with an area of 9.77 million km^2^ (7.31% of global land), are mainly located in the Sahara, southern Arabian Peninsula, and Greenland. The overlapping areas of the *A* region are rare, only accounting for 0.03% of terrestrial land.

#### 3.2.2. Protected area coverage

By overlapping the *A, B, C*, and *D* four regions with the global PA network, we found that in Africa and South America, many lands in the *D* region are covered by PAs (Fig. 6), while in terms of area, there are only a few PAs located in *the C* region. The PAs coverages in Europe countries are high, yet quite a proportion of those are located in *C* region besides in *D* region (Fig. S10 & S11).

**Fig. 6.**
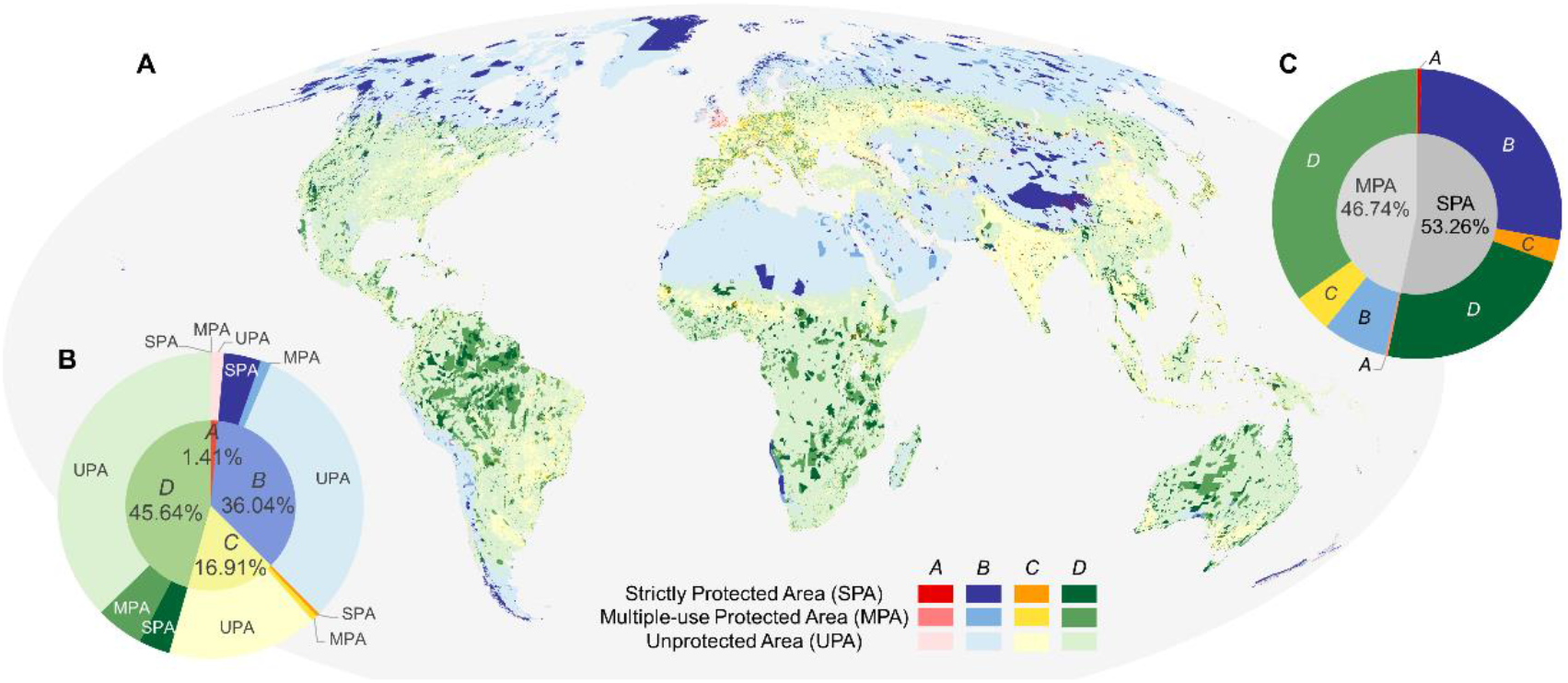
*Anthropic* (*A*), *Barren* (*B*), *Co-occurring* (*C*), and *Diversity-intact* (*D*) regions overlap with terrestrial protected area network. (a) Global map; (b) Hierarchical pie chart shows the percentage of the four regions worldwide (inner pie) and their overlaps with protected area (outer pie); (c) Hierarchical pie chart shows the proportion of terrestrial strictly protected area (SPA) and multiple-use protected area (MPA) worldwide (inner pie), and the proportion of the four regions within the two protected area categories (outer pie).

When the categories of PAs are taken into account, we found that 11.13 million km^2^ (18.27%) of the *D* region were covered by PAs, including 4.40 million km^2^ of strictly PAs and 6.73 million km^2^ of multiple-use PAs (Fig. 6a & 6b). Many *D* regions, including the western United States, the Amazon Basin, the Congo Basin, the Borneo, and central Australia, remain uncovered by PA network. The *C* region, where humans and wildlife co-occur, accounts for 16.91% of terrestrial land, with 0.49 million km^2^ (2.19%) and 0.83 million km^2^ (3.70%) of the *C* region covered by strictly and multiple-use PAs, respectively. While in the *B* region, which accounts for 36.04% of the global land, 5.28 million km^2^ (10.98%) of the *B* region lands are covered by strictly PAs and 1.39 million km^2^ (2.89%) by multiple-use PAs.

## 4. Discussion

By providing a series of spatial explicit indices, we reveal the worldwide land-sharing and land-sparing patterns between human and wildlife. Our findings indicate that the *D* region, with high biodiversity but relatively untouched by human activities, is the leading type (45.64%) on earth. Human activities are low in *D* and *B* regions, which are generally indiscriminately viewed as wilderness (Locke et al., 2019; Pimm et al., 2018). Our results, however, show that *B* and *D* regions differ significantly in their biodiversity. Most PAs distribute in *D* and *B* regions, yet the potential threats to the two regions are different. *D* region is vulnerable to direct anthropogenic impacts and its land use types are more prone to change, while the *B* region is often sensitive to global climate change (Fig. 2). Therefore, the objectives of conserved areas should be different in the two regions, and so do their management implementations. Surprisingly, lands merely dominated by humans (*A* region, 1.41%) are sparse, while the lands shared by humans and wildlife (*C* region, 16.91%) are far from insignificant since people never live independently from nature, even in the most densely populated region. Cities and farms are inhabited by a significant amount of wildlife, yield high values of the *C* index, and are of unignorable conservation value (Lepczyk et al., 2017; Li et al., 2020). Mace (Mace, 2014) suggested that the perspectives of conservation biology have shifted from ‘nature for itself’ to ‘people and nature’. The widespread *C* region confirms the necessity and urgency of enhancing conservation efforts and ecological research in those areas.

Our results suggest that different taxonomic groups might require different conservation strategies. Amphibians and reptiles have higher proportions of the *A* and *D* regions compared with their *C* regions, indicating a land-sparing strategy may necessary for them. Amphibians and reptiles are more sensitive to environmental changes and anthropogenic disturbances (Hopkins, 2007; Kafash et al., 2018; Russildi et al., 2016), so maintaining their habitats undisturbed from human activities play an important role for conservation. The *C* regions of birds and mammals are larger, especially, the *A* regions of amphibians and reptiles are often the *C* regions of birds and mammals. It implies that those two taxa can be protected through a land-sharing strategy. While it is necessary to strengthen conservation management toward human-wildlife relationships, such as mitigating human-wildlife conflict (Alexander et al., 2015; Struebig et al., 2018) and functional changes in species induced by human disturbance (Manlick and Pauli, 2020).

However, it is necessary to note that the species that benefit from land-sharing are often neither those of greatest conservation concern nor those found in the original native vegetation (Clough et al., 2011; Valente et al., 2022). For instance, in agroecosystems, where human and wildlife share lands, the composition of communities tend to be more homogeneous than that in the nearby forest due to habitat simplification (Valente et al., 2022). Despite the wide distribution and high proportion of the *C* regions in birds and mammals, the area and distribution of their *D* regions are more extensive. In many cases, the land-sparing area is essential for the protection of specialist species in specific ecosystems, as they are at higher risk of losing habitats than generalists (Baisero et al., 2020). Our results demonstrate that land-sharing and land-sparing are complementary conservation strategies to support diverse taxonomic groups and trade-off between human development and biodiversity conservation.

The *D* region is of high biodiversity and general conservation focus, essential for the global biodiversity conservation goals. Conservation actions should aim at establishing strictly-managed PAs in this region. However, our results reveal a mismatch between the current strictly PA network and the conservation needs. For instance, at least 11% (∼662,600 km^2^) of the natural rainforest in Amazonian was deforested in recent decades due to insufficient conservation (Bullock et al., 2020). Although Africa and South America have had large proportion of PAs coverage, they still hold tremendous potentials and promises for a land-sparing strategy, yet sufficient support is still required. Social and environmental justice should be considered and adequate resources must be secured before further expanding protected areas in those areas (Maxwell et al., 2020).

For a land-sharing strategy, the ways and effectiveness of establishing conserved areas in the *C* region will be quite different from those in the *D* region. The *C* region is the leading type in India, north China, and most European countries (Fig. 4). Given that the situations vary from case to case, it deserves the involvement and participation of indigenous peoples and other stakeholders, putting forward innovative ideas, backing sustainable development, and searching for compensation measures. In the *C* region, conservation management can be improved by multiple-use PAs, other effective area-based conservation measures or other practical approaches. Urban protected areas, adopted in different areas worldwide, have been proven valuable in creating wildlife habitats and providing essential ecosystem services (Ted Trzyna, 2014). If biodiversity can be effectively conserved in the *C* region, it would make the post-2020 global biodiversity framework more achievable.

The current patterns of HNIs are determined mainly by the expansion of human-dominated land-use. (Fig. 2). Since 1700, a large amount of natural vegetation has been changed to croplands (Ramankutty and Foley, 1999). Technological development and population growth have increased the scope of human activities on the earth’s surface. In the processes of human expansion, habitats have been losing and degrading, which in turn enormously impact global biodiversity (Newbold et al., 2016, 2015), leading to the formation of the current land-sharing and land-sparing patterns between human and wildlife, especially the large areas of *C* region in Eurasia. Global climates also shaped the pattern of HNIs. Mean temperature and annual precipitation are closely related to gross primary productivity (Zhang et al., 2017), thus driving the pattern of global biodiversity (Field et al., 2009; Gillman and Wright, 2013; Hawkins et al., 2003). Human distribution and the rise and fall of historical civilization have also been affected by climate (Haug et al., 2003; Small and Cohen, 2004). Such spatial patterns will be more complex and be persistent in the context of climate change, highlighting the necessity of limiting global warming to 1.5°C (Warren et al., 2018).

Our analysis brings a perspective in identifying priorities for future conservation efforts under the framework of land sharing and sparing on a global scale. Integrating human activity pressure, this set of HNIs advances our understanding on conservation priority solely focused on biodiversity and ecosystem services. The sharing-sparing patterns between human and wildlife vary considerably among countries and territories. This uneven distribution of the *A, B, C*, and *D* regions across countries indicates how each country contributes to the 30 by 30 goal should be differentiated. Sufficient thoughts and discussion would be beneficial before implementing the goals under the post-2020 global biodiversity framework. Our framework provides tools for future conservation implementations in different HNIs regions to achieve respective targets. We suggest that the dynamics of HNIs should be monitored and evaluated regularly. Besides, to achieve a more comprehensive understanding, it is also suitable and needed to evaluate the HNIs from other profiles, such as different taxa and levels of biodiversity or conservation status (Cai et al., 2020; Su et al., 2021).

## CRediT authorship contribution statement

Chengcheng Zhang: Conceptualization, Formal analysis, Writing – original draft.

Yihong Wang: Formal analysis, Writing – original draft.

Shengkai Pan: Formal analysis, Writing – original draft.

Biao Yang: Conceptualization.

Xiangjiang Zhan: Conceptualization, Funding acquisition.

Jiang Chang: Conceptualization, Funding acquisition.

Junsheng Li: Funding acquisition.

Qiang Dai: Conceptualization, Formal analysis, Writing – original draft, Funding acquisition.

## Data availability

Data will be made available on request.

## Declaration of competing interest

The authors declare no competing interests.

## Acknowledgements

This work was supported by the Strategic Priority Program of Chinese Academy of Sciences (grant number XDB31000000) and the National Natural Science Foundation of China (grant number 32070520).

## Notes

### Competing Interest Statement

The authors have declared no competing interest.

