## Supplementary Material for "Mapping global land sharing-sparing patterns between human and wildlife"

### Fig. S1.

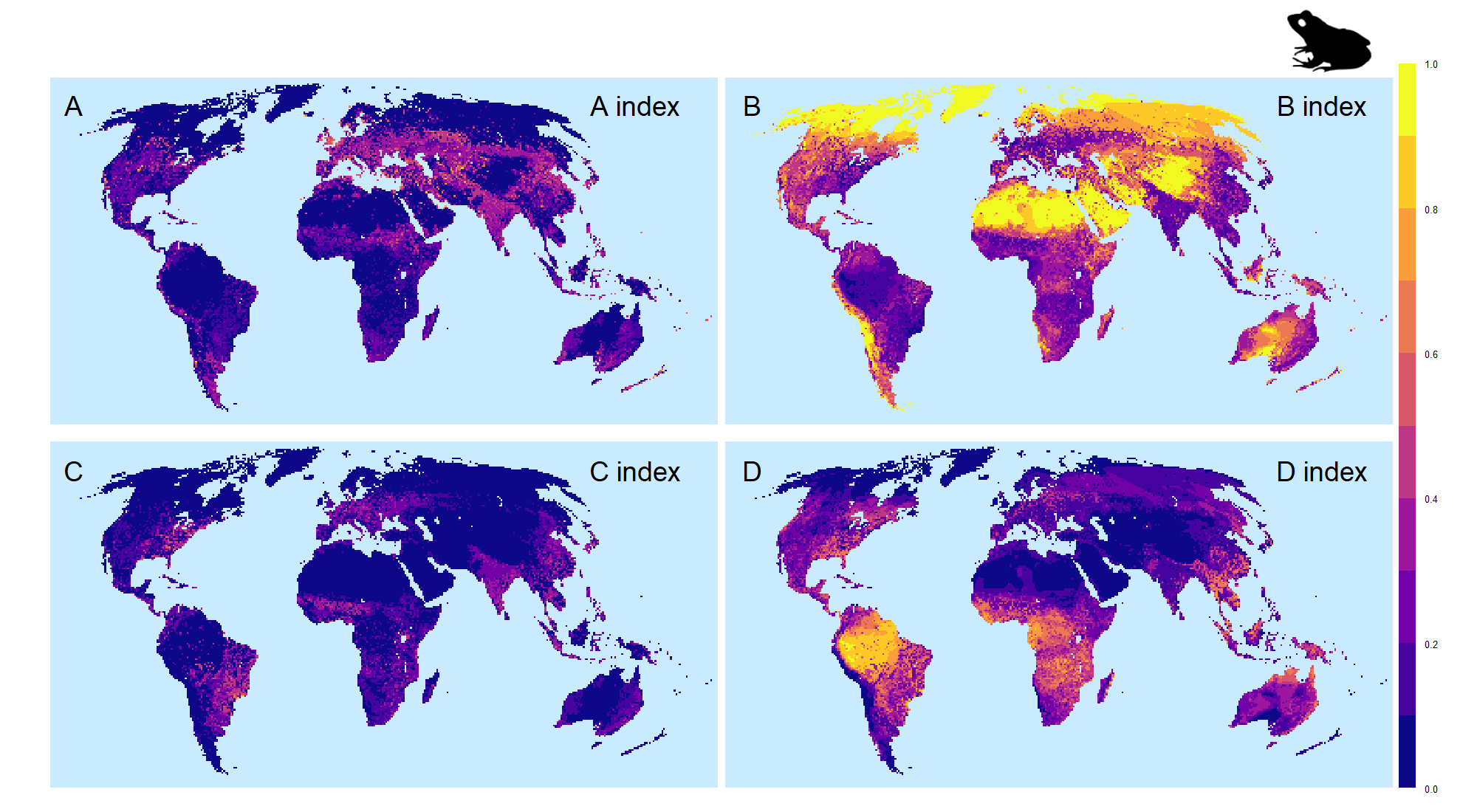

**Fig. S1.** Global land-sharing and land-sparing patterns between human and terrestrial amphibians illustrate by the four Human-Nature Indices (HNIs). (a) The *Anthropic* index (*A* index) indicates the extent that lands are dominated by humans; (b) the *Barren* index (*B* index) shows the extent that lands are neither occupied by amphibians nor by humans; (c) the *Co-occurring* index (*C* index) presents the extent of co-occurrence between human activities and amphibian species; and (d) the *Diversity-intact* index (*D* index) indicates the extent that lands are dominated by amphibians.

### Fig. S2.

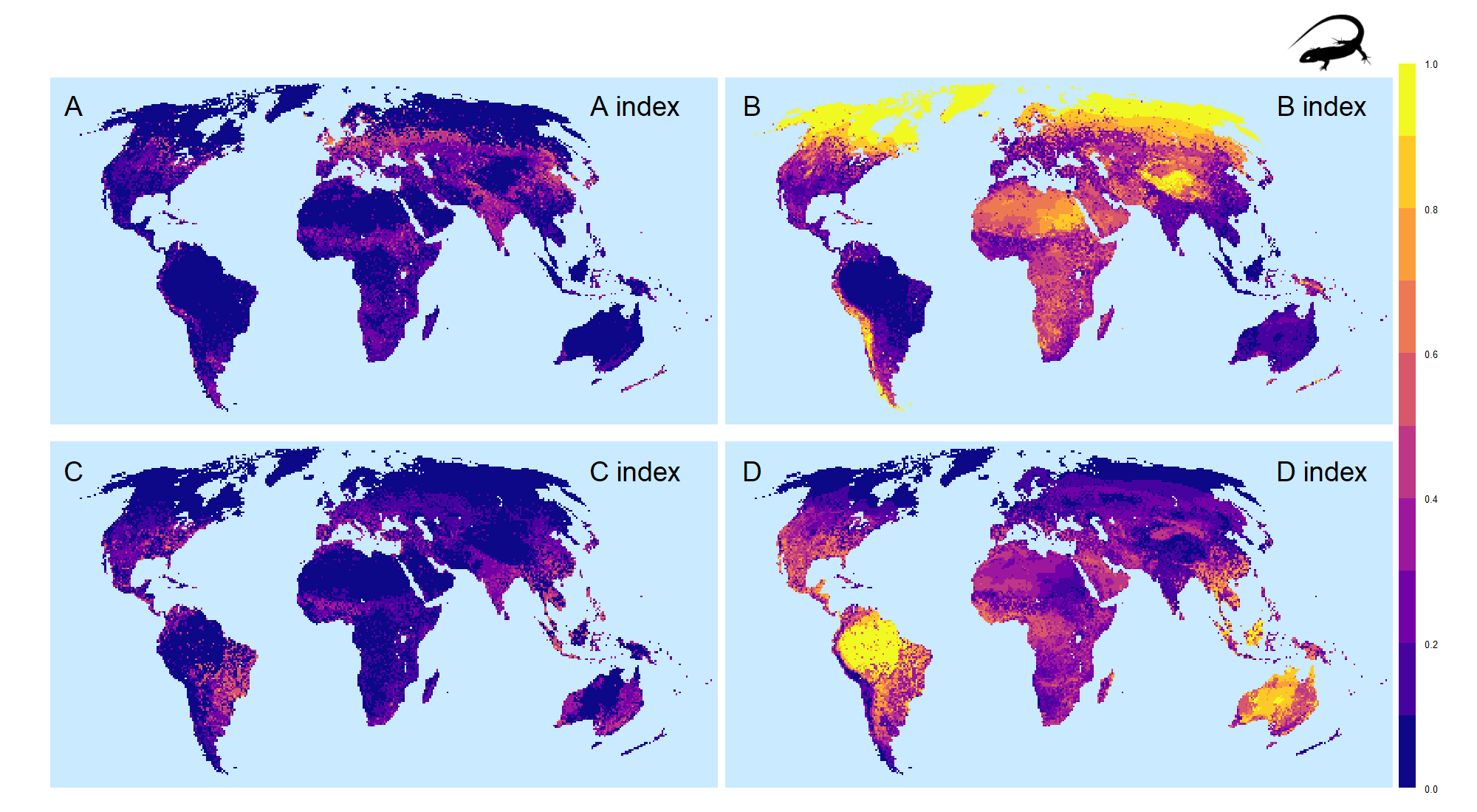

**Fig. S2.** Global land-sharing and land-sparing patterns between human and terrestrial reptiles illustrate by the four Human-Nature Indices (HNIs). (a) The *Anthropic* index (*A* index) indicates the extent that lands are dominated by humans; (b) the *Barren* index (*B* index) shows the extent that lands are neither occupied by reptiles nor by humans; (c) the *Co-occurring* index (*C* index) presents the extent of co-occurrence between human activities and reptile species; and (d) the *Diversity-intact* index (*D* index) indicates the extent that lands are dominated by reptiles.

### Fig. S3.

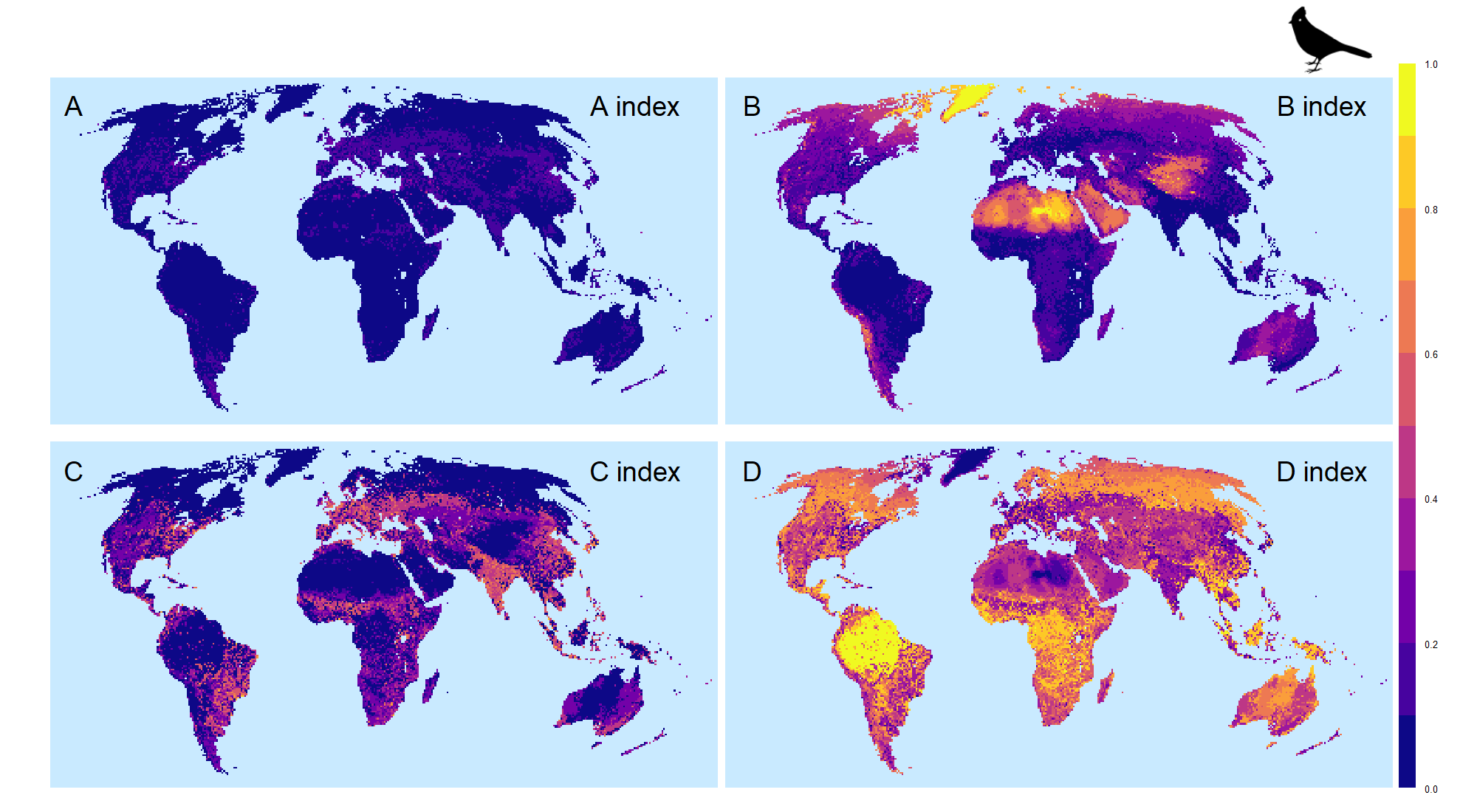

**Fig. S3.** Global land-sharing and land-sparing patterns between human and terrestrial birds illustrate by the four Human-Nature Indices (HNIs). (a) The *Anthropic* index (*A* index) indicates the extent that lands are dominated by humans; (b) the *Barren* index (*B* index) shows the extent that lands are neither occupied by birds nor by humans; (c) the *Co-occurring* index (*C* index) presents the extent of co-occurrence between human activities and bird species; and (d) the *Diversity-intact* index (*D* index) indicates the extent that lands are dominated by birds.

### Fig. S4.

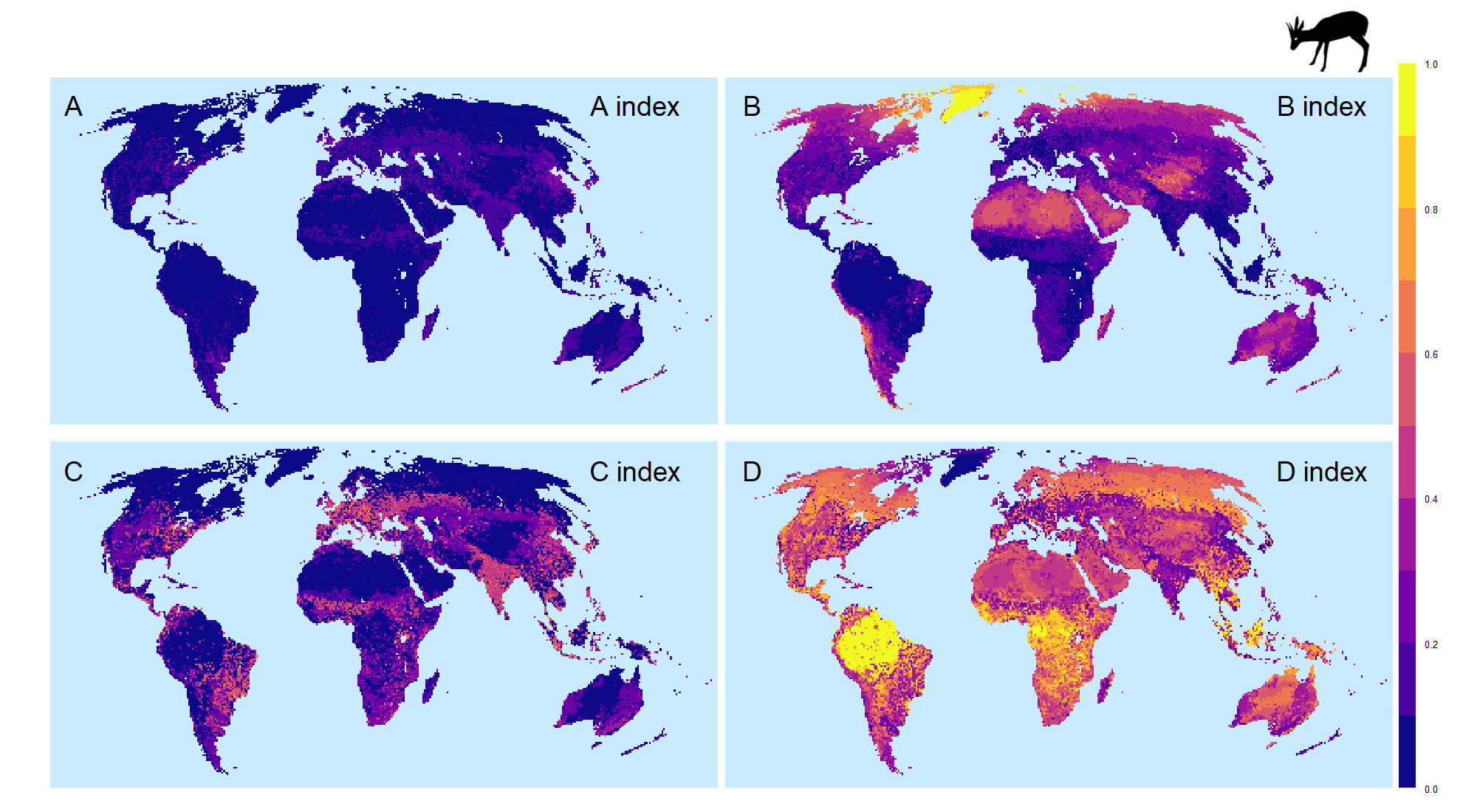

**Fig. S4.** Global land-sharing and land-sparing patterns between human and terrestrial mammals illustrate by the four Human-Nature Indices (HNIs). (a) The *Anthropic* index (*A* index) indicates the extent that lands are dominated by humans; (b) the *Barren* index (*B* index) shows the extent that lands are neither occupied by mammals nor by humans; (c) the *Co-occurring* index (*C* index) presents the extent of co-occurrence between human activities and mammal species; and (d) the *Diversity-intact* index (*D* index) indicates the extent that lands are dominated by mammals.

### Fig. S5.

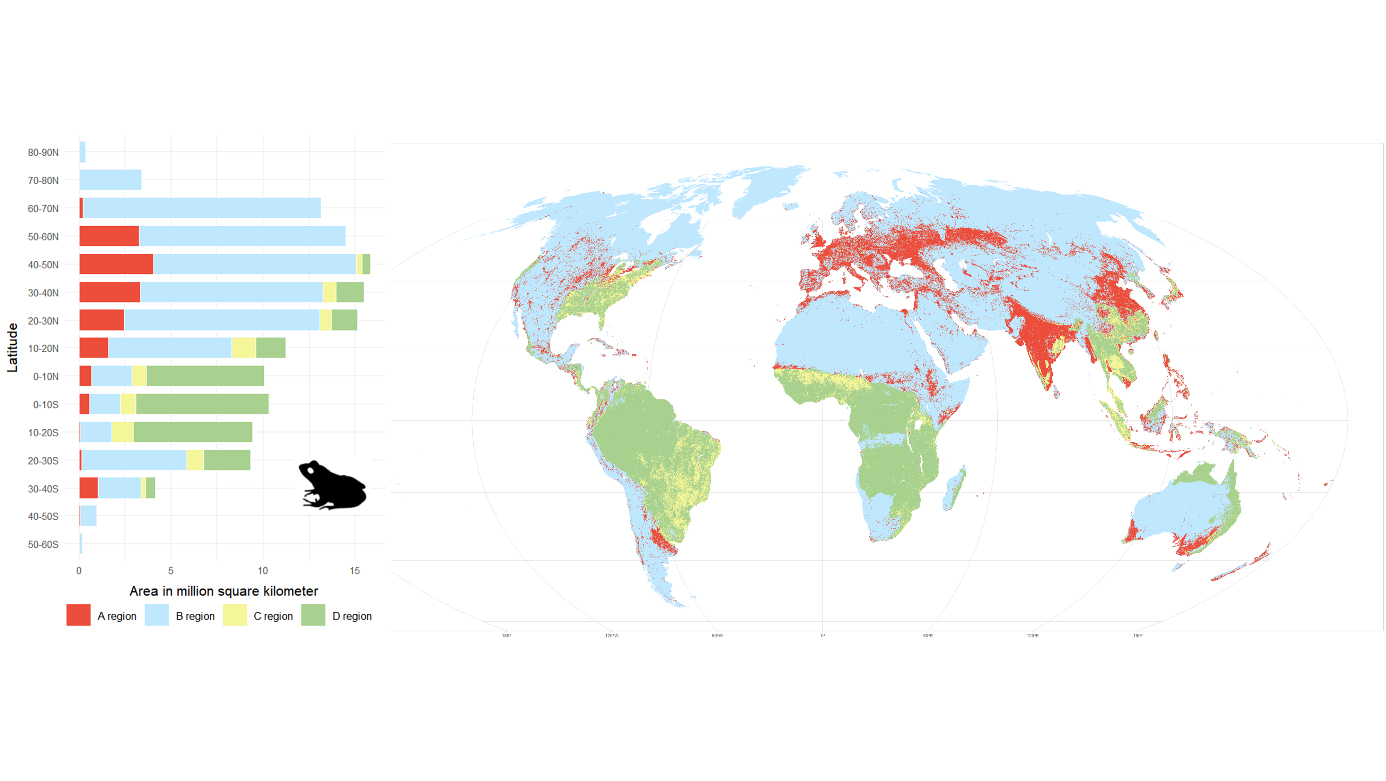

**Fig. S5.** *Anthropic* (*A*), *Barren* (*B*), *Co-occurring* (*C*), and *Diversity-intact* (*D*) regions illustrate the land-sharing and land-sparing statues between human and terrestrial amphibians. Red indicates the *A* region, blue indicates the *B* region, yellow indicates the *C* region, and green indicates the *D* region.

### Fig. S6.

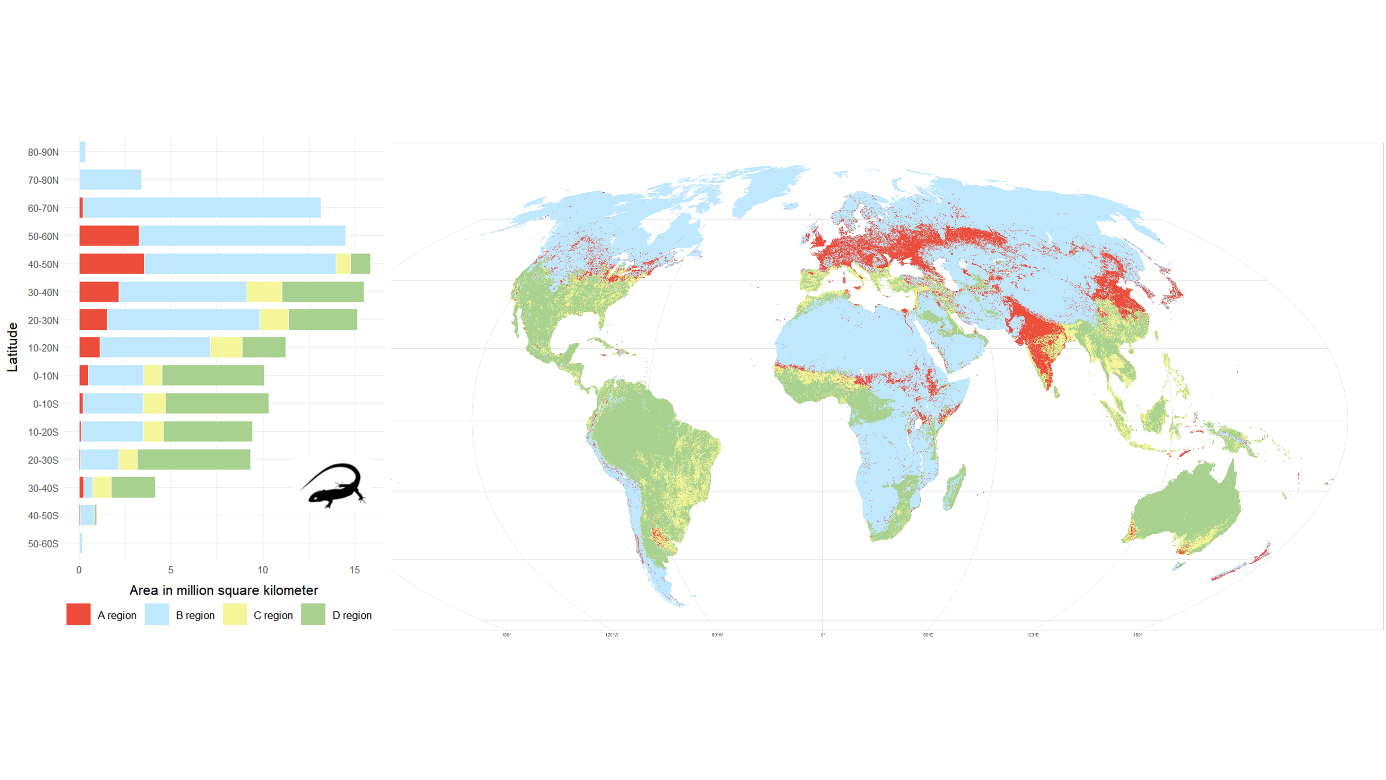

**Fig. S6.** *Anthropic* (*A*), *Barren* (*B*), *Co-occurring* (*C*), and *Diversity-intact* (*D*) regions illustrate the land-sharing and land-sparing statues between human and terrestrial reptiles. Red indicates the *A* region, blue indicates the *B* region, yellow indicates the *C* region, and green indicates the *D* region.

### Fig. S7.

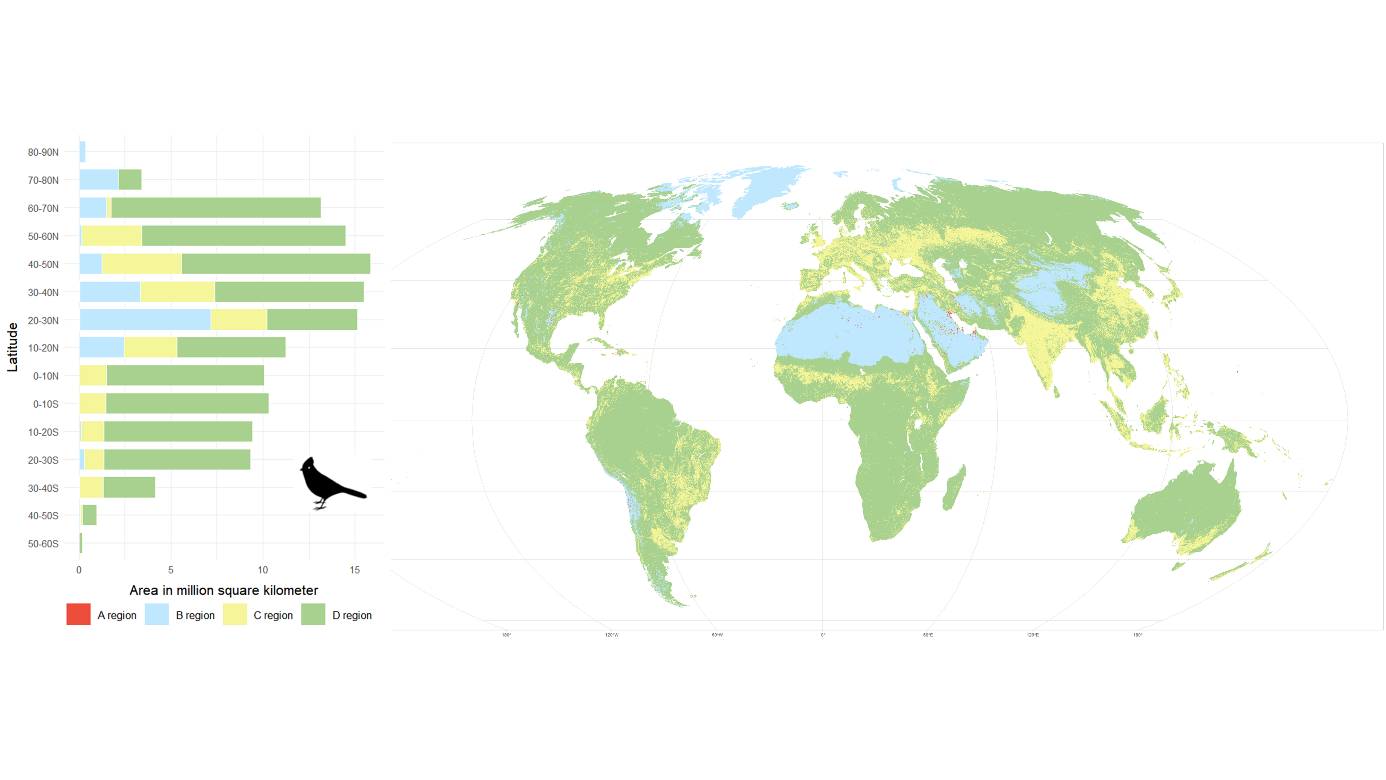

**Fig. S7.** *Anthropic* (*A*), *Barren* (*B*), *Co-occurring* (*C*), and *Diversity-intact* (*D*) regions illustrate the land-sharing and land-sparing statues between human and terrestrial birds. Red indicates the *A* region, blue indicates the *B* region, yellow indicates the *C* region, and green indicates the *D* region.

### Fig. S8.

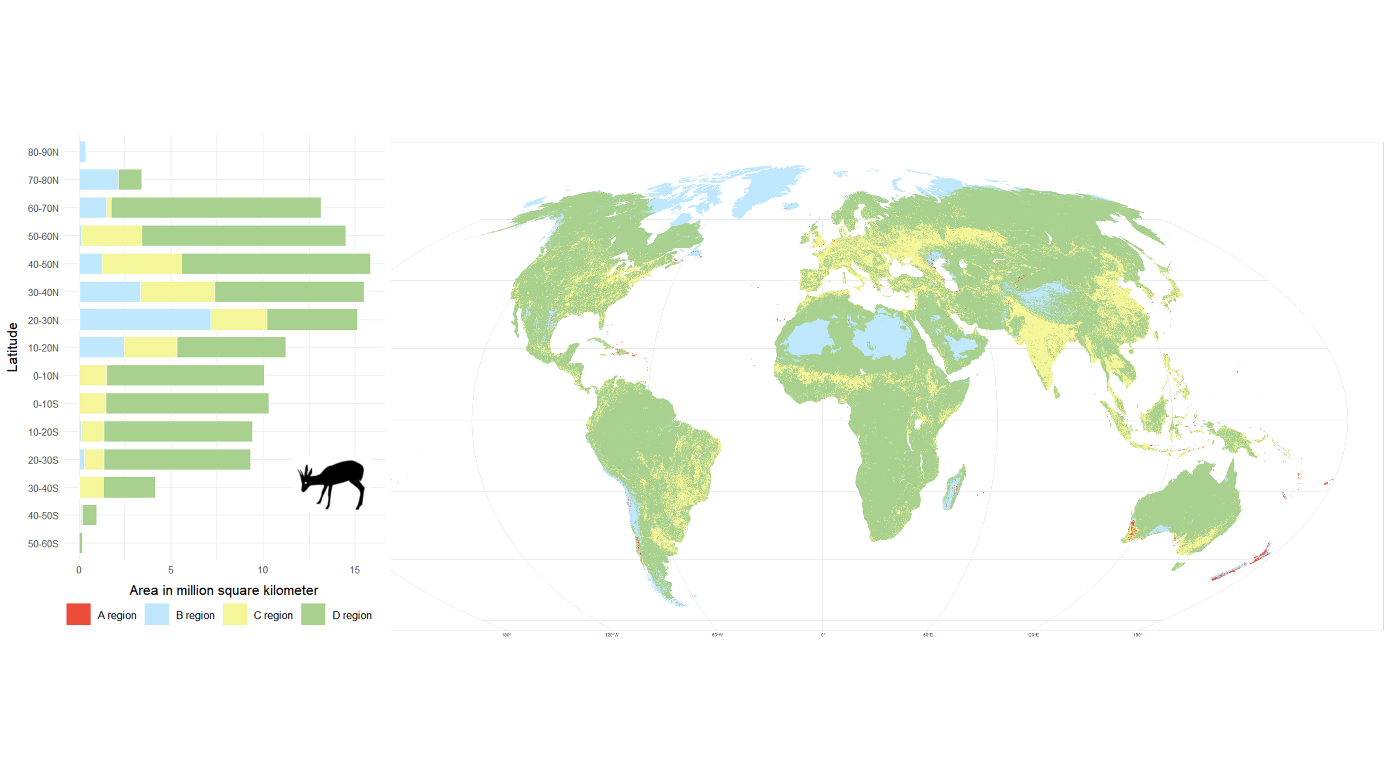

**Fig. S8.** *Anthropic* (*A*), *Barren* (*B*), *Co-occurring* (*C*), and *Diversity-intact* (*D*) regions illustrate the land-sharing and land-sparing statues between human and terrestrial mammals. Red indicates the *A* region, blue indicates the *B* region, yellow indicates the *C* region, and green indicates the *D* region.

### Fig. S9.

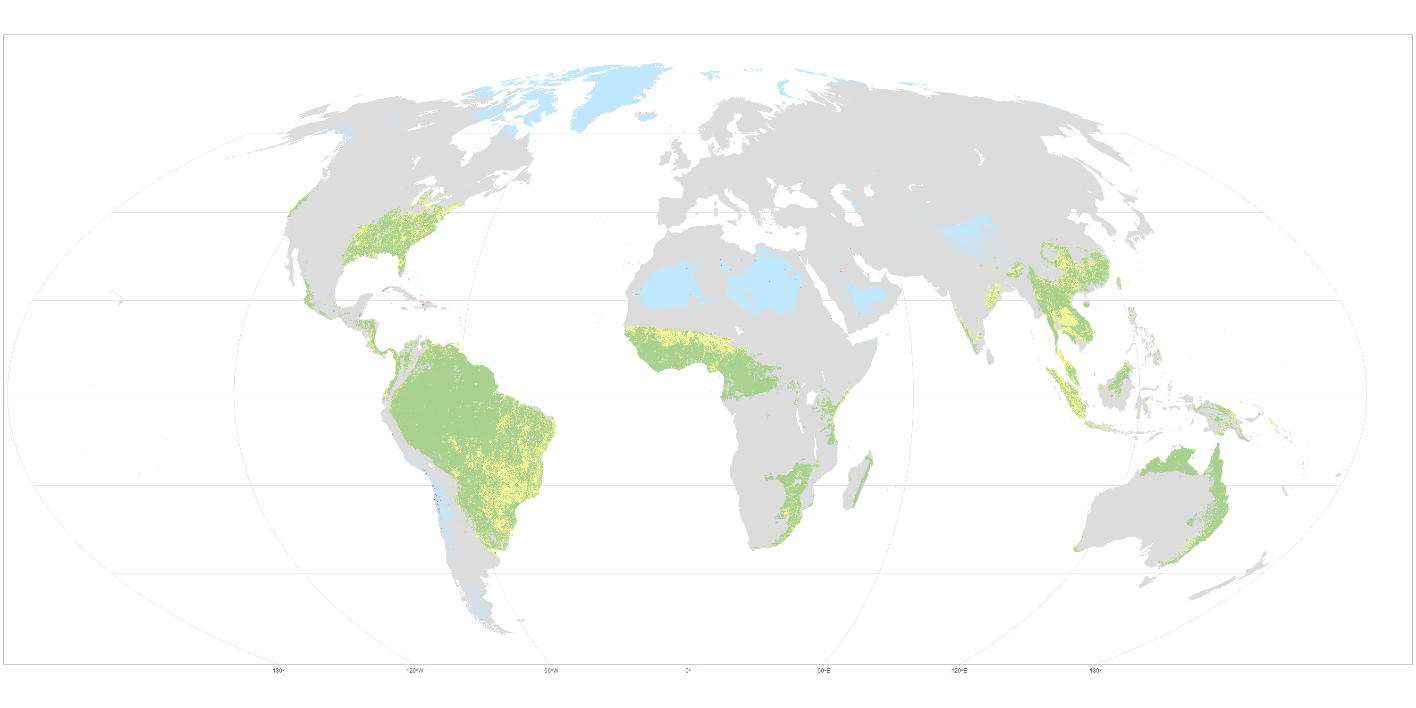

**Fig. S9.** The overlaps of the *Anthropic* (*A*), *Barren* (*B*), *Co-occurring* (*C*), and *Diversity-intact* (*D*) regions of the four taxonomic groups (amphibians, reptiles, birds and mammals). Red indicates the overlaps of the *A* regions, blue indicates the overlaps of the *B* regions, yellow indicates the overlaps of the *C* region, and green indicates the overlaps of the *D* region.
